# The Rod Bipolar Cell Pathway Contributes To Surround Responses In OFF Retinal Ganglion Cells

**DOI:** 10.1101/2025.10.01.679777

**Authors:** G. Spampinato, T. Trapani, V. Calbiague-Garcia, T. Buffet, E. Orendroff, B.S. Sermet, D. Dalkara, E. Ronzitti, E. Papagiakoumou, V. Emiliani, O. Marre

## Abstract

Sensory neurons can be influenced by stimuli beyond their receptive field center, yet the mechanisms underlying this surround modulation remain poorly understood. In the retina, many OFF ganglion cells exhibit responses to ON stimulation outside their receptive field center. However, disentangling the pathways and cell types contributing to these responses has been challenging with traditional experimental approaches. Here, we combined optogenetics, two-photon holographic stimulation, and multi-electrode array recordings to identify the intermediate retinal cell types involved in this circuit. We found that the pathway formed by rod bipolar cells and AII amacrine cells—one of the primary relays of rod-driven signals under low-light conditions—plays a key role in mediating this surround modulation. Specifically, crossover inhibition exploits the same amacrine cells responsible for surround suppression to disinhibit distant ganglion cells. This suggests that the retina repurposes existing circuits for surround modulation, optimizing resources through multifunctional inhibitory pathways.

## Introduction

A major feature of sensory neurons is that they can perform complex computations with limited resources. Understanding how they achieve this feat is a major challenge for sensory neuroscience. This challenge arises because, rather than being implemented by a single dedicated pathway, several circuits often contribute to the computations performed by a sensory neuron, each participating in different facets of this computation.

Surround modulation is a paramount example of such computation. It is a well-established phenomenon in the retina and other neural circuits, where stimuli presented outside the receptive field center influence neuronal responses. For example, when displaying a large disc centered on a retinal ganglion cell, the stimulation of the surround suppresses the responses to the center (Kuffler, 1953; Farrow et al., 2013; Thoreson and Mangel, 2012). As a result, many ganglion cells respond maximally to a stimulus of optimal size. However, this surround suppression is not the only feature of surround modulation. Many ganglion cells can also be modulated by stimuli outside their receptive field center, with an opposite polarity to their preferred one: a ganglion cell selective to OFF stimuli in its center may respond to ON stimuli in the surround (Bloomfield and Xin, 2000; Kamermans and Spekreijse, 1999). This phenomenon will be termed antagonistic surround modulation in the following.

It is unclear if these different facets of surround modulation are generated by the same mechanism or by different ones. Horizontal cells are traditionally thought to play a central role in surround suppression by mediating lateral inhibition through their synapses with photoreceptors (Davenport et al., 2008; Mangel, 1991; McMahon et al., 2004; Werblin, 1972). However, other studies suggest alternative or additional mechanisms where amacrine cells are also involved in surround suppression, providing inhibitory inputs that shape the response properties of ganglion cells, either directly inhibiting ganglion cells or indirectly by inhibiting the bipolar cell input (Cook and McReynolds, 1998; Farrow et al., 2013; Franke et al., 2017; Jensen, 1991; Johnson et al., 2018; Nath et al., 2023; Park et al., 2020).

Recent evidence suggests that rods may also contribute to surround modulation, especially at mesopic light levels. Rods remain active over a broader range of light intensities than previously believed and can influence cone-mediated pathways (Grimes et al., 2018; Pang et al., 2003; Tikidji-Hamburyan et al., 2015). Rods can contribute to surround modulation through horizontal cells, which laterally inhibit cone output, thereby modulating bipolar cell activity.

Another pathway that may contribute to surround modulation is the rod bipolar cell (RBC)-AII amacrine cell circuit, which has been shown to remain active at intermediate light levels (Franke et al., 2017; Ke et al., 2014). Anatomical studies have suggested a disinhibitory pathway involving RBCs, AII amacrine cells, OFF cone bipolar cells, wide-field GABAergic amacrine cells, and OFF ganglion cells (Fig. 5) (Lauritzen et al., 2019), through which rod bipolar cells could modulate OFF ganglion cell responses, but this has not been confirmed experimentally so far.

This raises the question of whether RBC or AII amacrine cell stimulation can participate in antagonistic surround modulation by affecting the activity of distant OFF ganglion cells. Consistent with this hypothesis, it has been shown that the responses of OFF ganglion cells to ON stimulation in the surround depend on a glycinergic pathway, as responses are blocked by strychnine (Deny et al., 2017; Franke et al., 2017). However, existing experimental approaches lack the specificity needed to precisely establish whether the RBC-AII pathway directly contributes to these antagonistic responses. For instance, strychnine blocks all glycinergic pathways, and rod activation can engage additional circuits, such as those mediated by horizontal cells.

To address this question, we employed a novel combination of 2-photon computer-generated holography, optogenetics, and multi-electrode recordings, enabling precise spatial and cell-type-specific stimulation. This approach allowed us to examine whether the RBC-AII pathway contributes directly to the antagonistic surround modulation observed in OFF ganglion cells. We tested whether activating the RBC-AII pathway alone is sufficient to evoke responses in distant OFF ganglion cells, effectively mimicking the antagonistic ON surround responses. Additionally, we found that hyperpolarizing AII amacrine cells significantly reduces the responses of OFF ganglion cells to ON surround stimulation. Together, these findings show that the RBC-AII pathway contributes to antagonistic surround modulation in OFF ganglion cells. Beyond highlighting a new role for the AII amacrine cells, this suggests that complementary mechanisms mediate surround suppression and antagonistic surround modulation: ON stimulation will evoke crossover inhibition of the OFF pathway. This inhibition will inhibit the wide-field OFF amacrine cells that mediate surround suppression and disinhibit OFF ganglion cells. Crossover inhibition thus “piggybacks” on the amacrine cells mediating surround suppression to implement antagonistic surround modulation. This shows how a neural circuit can be recycled in different contexts for different computational roles.

## Results

### RBCs Activate OFF Ganglion Cells Beyond Their Receptive Field Center

To test whether the RBC-AII pathway could evoke responses in distant OFF ganglion cells, we first stimulated single RBCs. Our previous work has shown that 2-photon holographic stimulation allows for evoking depolarization of RBCs that mimic the kind of depolarization evoked by light stimuli (Spampinato et al., 2022). We developed a tool combining holographic stimulation at 1040 nm with multi-electrode array (MEA) recordings (Yger et al., 2018), which allowed us to stimulate RBCs with high spatial precision while recording ganglion cell activity (Fig. 1A). This system allowed generating holographic spots with a 10 µm diameter in the x-y plane—matching the diameter of RBCs—and a 22 µm resolution along the z-axis, slightly large but still smaller than the full z-axis extent of an RBC from dendrite to axon terminal (Fig. 1B, C).

**Fig. 1.**
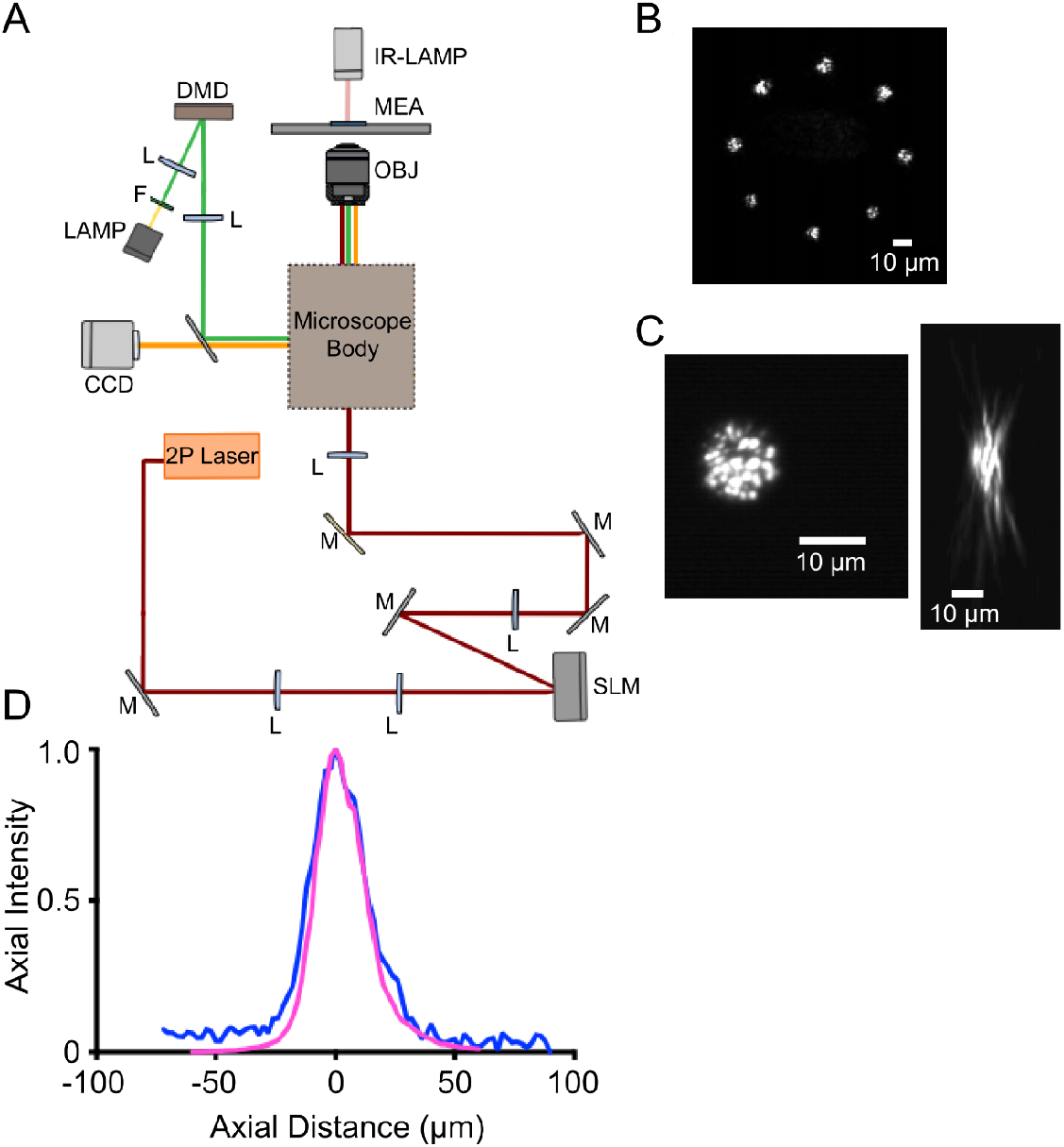
Setup description and characterization. **A**. Schematic of the optical system. All the optical paths are shown in different colors. F: filter, L: lens, M: mirror. **B**. Image of 8 spots at the borders of the area we used for the photostimulation. Laser intensity is roughly homogeneous inside this area. **C**. Lateral and axial profile of a 10 µm holographic spot. Fluorescence image excited at 1040 nm by a 10-µm-diameter spot on a rhodamine thin layer. **D**. Axial profile of the integrated fluorescence intensity of a 10 µm holographic spot. The Magenta curve is the measure without the MEA, and the blue curve is the same measure through the MEA glass.

To make RBCs light-sensitive, we expressed the optogenetic protein CoChR fused with GFP specifically in these cells. This was achieved by injecting an AAV vector (Dalkara et al., 2013) under the control of a cell-type-specific promoter (Lu et al., 2016), resulting in restricted expression in RBCs, similar to prior work (Fig. 2B) (Spampinato et al., 2022).

**Fig. 2.**
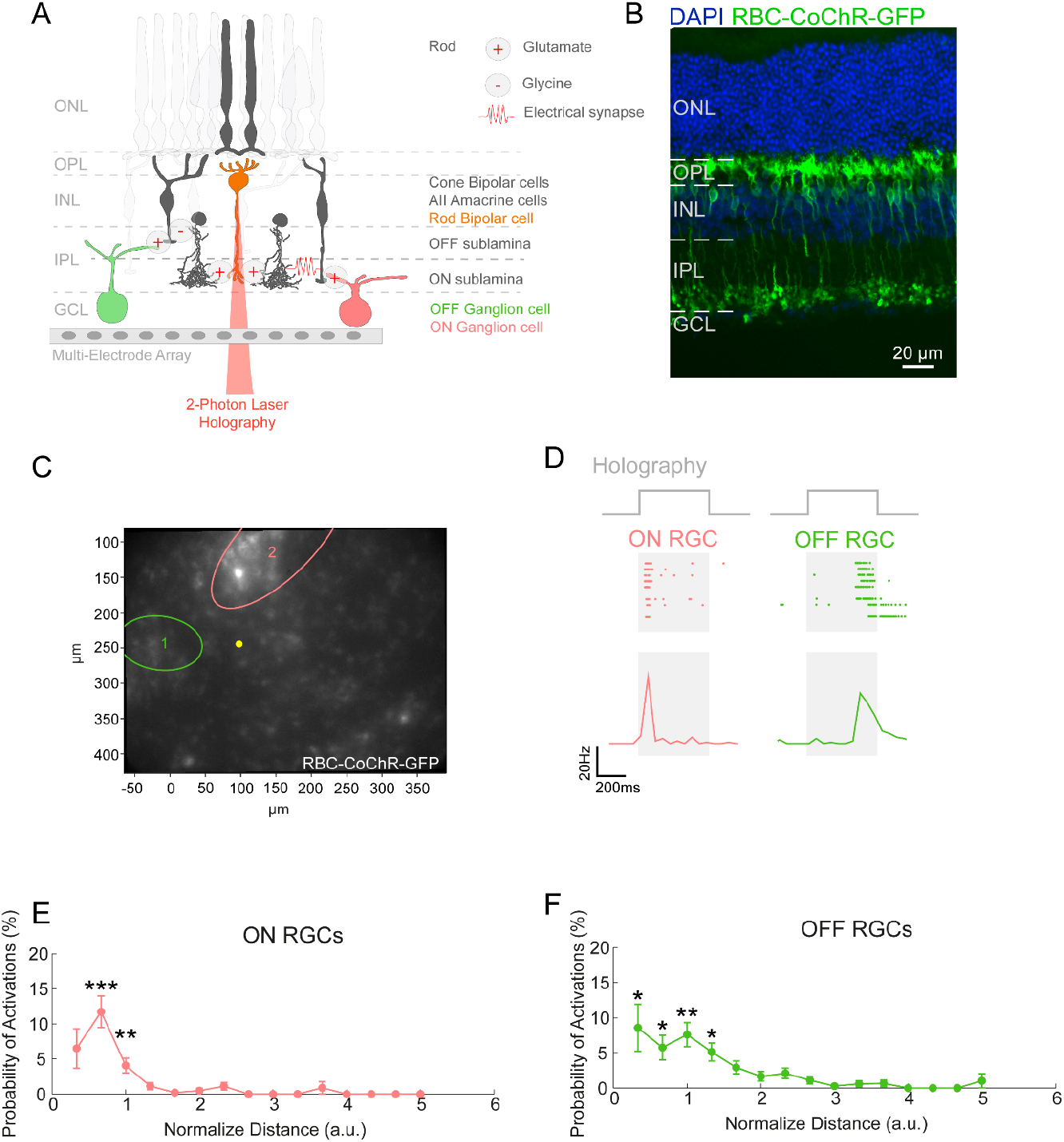
Optogenetic stimulation of RBCs evokes ganglion cell responses. **A**. Schematic of the experimental protocol. Rod bipolar cells (RBCs) are optogenetically stimulated, activating a polysynaptic pathway involving AII amacrine and cone bipolar cells. Retinal ganglion cells integrate this signal, generating responses recorded using multi-electrode arrays (MEA). **B**. Retinal slice showing the expression of CoChR-GFP in rod bipolar cells. **C**. Image of the bipolar cell layer during the experiment, with RBCs identified by their fluorescence halo. A representative RBC targeted by optogenetic stimulation is highlighted in yellow. The receptive fields (RFs) of two representative ganglion cells (one ON and one OFF) are overlaid in red and green, respectively. **D**. Responses of the ON (red) and OFF (green) ganglion cells shown in C to RBC activation. **Top:** Spiking activity across trials. **Bottom:** Mean responses of the two ganglion cells. The gray regions indicate the stimulation period. **E**. Probability of an ON ganglion cell response to RBC stimulation as a function of the relative distance between the ganglion cell and RBC, with distances normalized by the ganglion cell RF center radius. **F**. Same as D for OFF ganglion cells. Graphs display mean values ± standard error; asterisks indicate *p < 0.05, **p < 0.01, ***p < 0.001.

Then, we conducted the following experiment: (1) We recorded ganglion cell activity during visual stimulation. (2) We mapped each ganglion cell’s receptive field (RF) using a checkerboard stimulus. (3) We blocked photoreceptor-to-bipolar cell transmission with a pharmacological cocktail (15 µM L-AP4; 10 µM ACET). (4) Using epifluorescence microscopy, we identified RBCs expressing CoChR-GFP. (5) Finally, we stimulated individual RBCs while recording ganglion cell activity (Fig. 2A).

Stimulation of RBCs elicited responses in ganglion cells. We found that OFF ganglion cells responded to distant stimulation of RBC, even when the stimulated RBC was located outside the ganglion cell’s receptive field center (Fig. 2C, D). To quantify these responses, we analyzed all RBC-ganglion cell pairs and identified significant responses (defined as a z-score above a threshold and a firing rate above a minimum value; see Methods). RBC stimulation evoked responses in both ON and OFF ganglion cells. To assess the relationship between response probability and distance, we plotted the probability of evoked responses against the normalized distance between the stimulated RBC and the RF center of the recorded ganglion cell (normalized such that 1 corresponds to the RF center’s boundary). For both ON and OFF ganglion cells, the estimated probability of observing a response when stimulating an RBC in their receptive field center was ∼7% (Fig. 2E, F). For ON ganglion cells, the probability of a response declined rapidly beyond the RF center boundary (< 1% for normalized distances between 1.00 and 1.33, *N* = 2 wild-type retinas, *n* = 93 ganglion cells, *p* = 0.1295, Fig. 2E). In contrast, OFF ganglion cells responded to RBC stimulation at greater distances, clearly outside of their RF center (∼5% for normalized distances between 1.00 and 1.33, *N* = 2 wild-type retinas, *n* = 118 ganglion cells, *p* = 0.0247, Fig. 2F). These results demonstrate that RBC stimulation can evoke responses in ganglion cells, including those located outside the RF center, as in the case for OFF ganglion cells.

### Ganglion cells Responses Cannot Be Explained By Ineffective Blocking Of Photoreceptor Transmission

We then controlled if the observed responses could be due to holographic stimulation of photoreceptors. Although holographic stimulation was performed using infrared light, it could still activate photoreceptors. We performed control experiments where we did the same protocol in mice that were not injected with AAV and did not express CoChR. Without applying pharmacological blockers, ganglion cells showed clear responses to holographic stimulation. There was a strong correlation between holographic stimulation and classical visual stimulation: holographic spots placed near the receptive field center elicited an increase in firing rate for ON ganglion cells and a decrease for OFF ganglion cells (*N* = 2 wild-type retinas, *n* = 110 ganglion cells, Fig. S1A and B). This opposite modulation for ON and OFF cells suggests that these responses to holographic stimulation are due to photoreceptor activation and are unlikely due to direct stimulation of the ganglion cell.

We then applied the same pharmacological cocktail we used previously to block the transmission from photoreceptors to bipolar cells. The responses to holographic stimulation entirely disappeared (Fig. S1A and C; see Methods). In some cases, however, we observed a slight and constant decrease in firing rate during holographic stimulation (Fig. S1A). Previous studies (Owen et al., 2019; Picot et al., 2018) have shown that two-photon stimulation can locally increase temperature, leading to potassium channel activation and a slight hyperpolarization. Our findings are consistent with this mechanism, suggesting that this small and constant suppression is likely due to local heating of ganglion cells caused by holographic stimulation. The sustained nature of this suppression further supports this hypothesis. These effects were clearly different from the responses observed above when stimulating rod bipolar cells, demonstrating that our results were not due to an incomplete blockade of photoreceptor transmission.

### Glycinergic transmission contributes to the surround responses of OFF ganglion cells

Our findings demonstrate that depolarizing RBCs with optogenetics can evoke responses in distant OFF ganglion cells. Thus, they may contribute to a circuit that enables OFF ganglion cells to respond to distant ON visual stimuli (antagonistic surround responses). This hypothesis is supported by the fact that RBCs can respond to ON stimuli even at mesopic light levels (Franke et al., 2017; Ke et al., 2014; Pang et al., 2003). AII amacrine cells, the primary recipients of RBC input, modulate the OFF pathway through their glycinergic synapses onto OFF bipolar cells. We thus hypothesized that the RBC-AII pathway plays a key role in mediating the ON antagonistic surround modulation observed in OFF ganglion cells.

To determine whether glycinergic transmission contributes to the antagonistic surround, we stimulated OFF ganglion cells with a distant bright disc positioned outside their receptive field center. After applying strychnine (10 µM) to block glycinergic transmission, the responses to the distant bright disc disappeared (*N* = 3 retinas, *n* = 35 ganglion cells, p < 0.001, Fig. 3). This indicates that antagonistic surround responses rely on glycinergic transmission, confirming previous findings (Cook and McReynolds, 1998; Deny et al., 2017; Franke et al., 2017). However, the retina contains multiple glycinergic amacrine cell types beyond AII amacrine cells, and this experiment does not demonstrate that AII amacrine cells are involved in the antagonistic surround modulation of OFF ganglion cells.

**Fig. 3.**
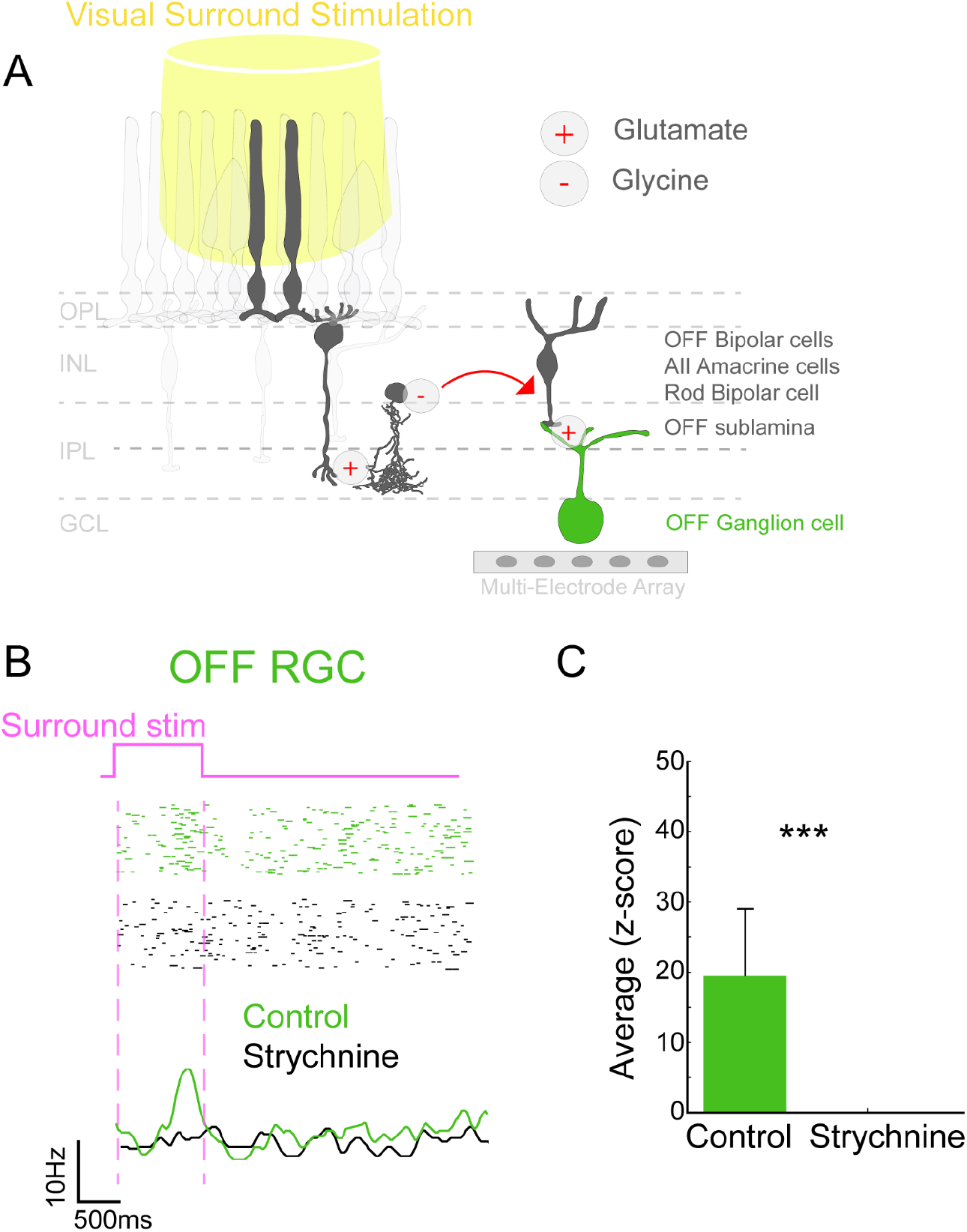
Glycinergic amacrine cells contribute to the surround responses of OFF ganglion cells. **A**. Schematic of the experimental protocol: a disc of white light (represented by the yellow cylinder) is flashed above the retinal tissue. The responses produced by the OFF ganglion cells are recorded with a multi-electrode array. **B**. Surround responses of a representative OFF ganglion cell. Top, spiking activity of the OFF ganglion cell to the visual white disc stimulus in the absence of a blocker (black). Middle, spiking activity of the same OFF ganglion cell, after the application of the glycinergic blocker strychnine (red). Bottom, the mean responses in both conditions. The violet dashed-line area represents the time interval of the surround stimulation. **C**. Quantification of the responses triggered by the surround stimulation with and without strychnine. Graphs display mean values ± SD; asterisks indicate ***p < 0.001.

### Hyperpolarizing The RBC-AII Pathway Affects Responses Of OFF Ganglion Cells To Surround Stimulation

To test whether AII amacrine cells are involved in antagonistic surround response of OFF ganglion cells, we modulated them directly. For this, we designed an experiment to inactivate the AII amacrine cells while simultaneously delivering visual and holographic stimulation (Fig. 4A) to test its contribution to ON surround responses. This approach allowed us to modulate the RBC-AII pathway during visual stimulation.

**Fig. 4.**
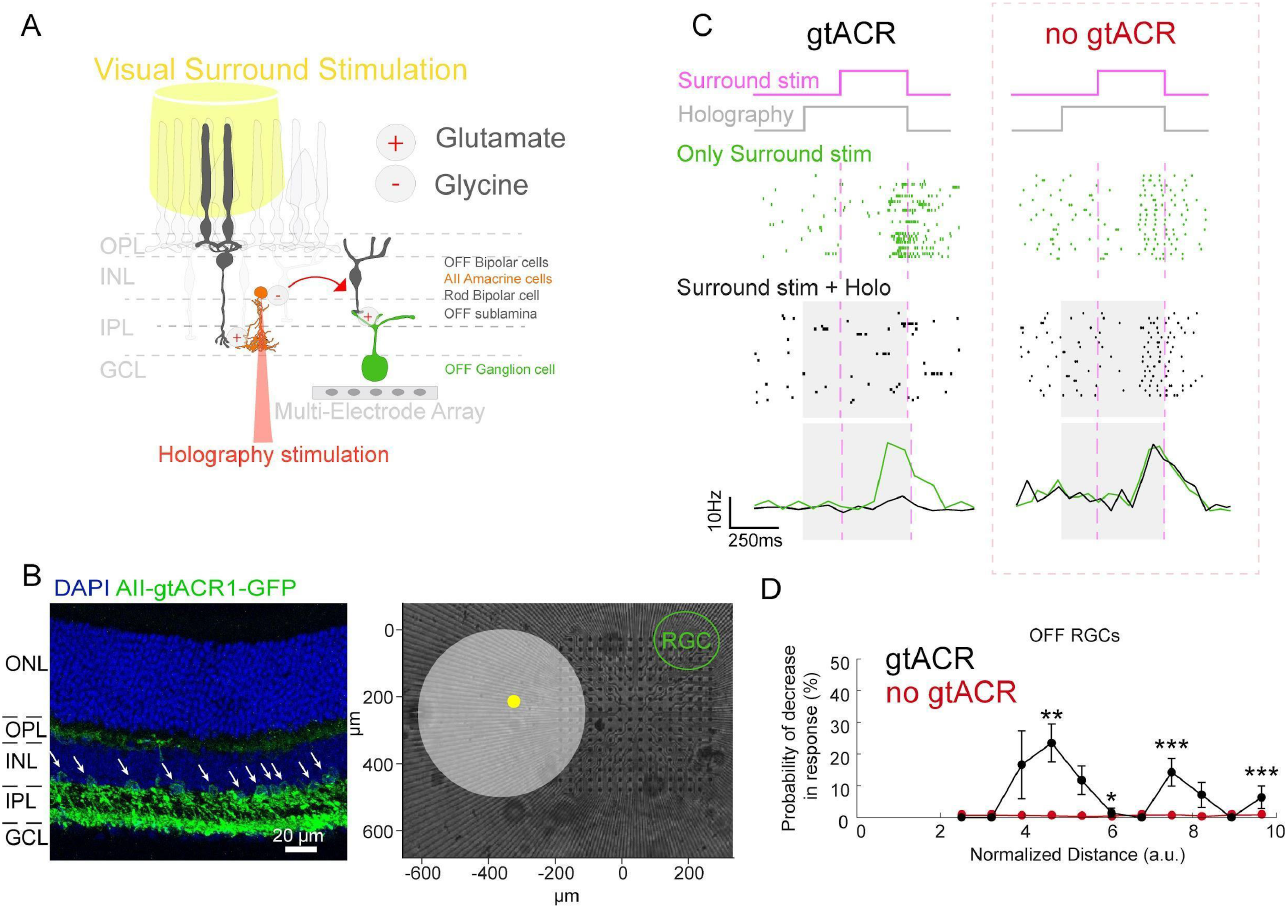
AII Amacrine cells contribute to the surround responses of OFF ganglion cells. **A** Schematic of the experimental protocol: a disc of white light is flashed above the retinal tissue (yellow). Simultaneously, a holographic spot inside the disc’s perimeter inhibits the underlying AII amacrine cells. The responses produced by the ganglion cells are recorded with a multi-electrode array. **B. (Left)** Retinal slice showing the expression of gtACR1-GFP in AII amacrine cells. **(Right)** relative positions of the disc (in white), a representative holographic spot (in yellow), the receptive field center of an OFF ganglion cell (in green), and the multi-electrode array (image in the background). **C. (left)** Surround responses of the representative OFF ganglion cell shown in B. Top, spiking activity of the OFF ganglion cell to the purely visual white disc stimulus (green). Middle, spiking activity to the combined visual and holographic stimulus (black). Bottom, the mean responses. The grey-shaded area represents the time interval of the holographic stimulus. The two violet dashed lines show the onset and offset of the visual stimulus. **(right)** Same as C for a retina in the absence of opsin, showing no significant modulations on OFF ganglion cells’ surround responses. **(D)** Probability of observing a decrement (of 10 Hz or more) in OFF ganglion cell surround responses when AIIs are inhibited, against the relative distance between the ganglion cell and the location targeted by the inhibitory stimulus. Distances were normalized by dividing by the radius of the ganglion cell receptive field center. Graphs display mean values ± standard error; asterisks indicate *p < 0.05, **p < 0.01, ***p < 0.001.

For this purpose, we expressed the hyperpolarizing-optogenetic protein gtACR1 in AII amacrine cells (Fig. 4B) (Govorunova et al., 2015), using an AAV construct with a promoter specific to these amacrine cells, which we tested previously (Khabou et al., 2023). We then designed an experiment where an ON visual stimulus was presented in the surround of OFF ganglion cells, with and without simultaneous holographic stimulation targeting AII amacrine cells. We hypothesize that AII amacrine cells should relay the signal from RBCs activated by ON surround stimulation to ganglion cells through a polysynaptic pathway (Fig. 4A). Therefore, hyperpolarizing AII amacrine cells should impair this transmission and reduce the responses of OFF ganglion cells to ON surround stimulation. The visual stimulus was as shown in Figure 3: a bright disc presented in the surround of recorded OFF ganglion cells. For holographic stimulation, one spot within the visual disc was targeted (Fig. 4B).

A technical challenge we faced was that holographic stimulation also activates photoreceptors (Fig. S1). We expected possible ON responses from photoreceptor activation by holographic stimulation for OFF ganglion cells stimulated in their surround. To separate the effect of holographic stimulation of photoreceptors from the effect of holographic stimulation of the AII amacrine cells, we introduced a delay between the holographic stimulation and the visual stimulation. Holographic stimulation of a single spot lasted 500 ms, starting 175 ms before the onset of the visual stimulus, which lasted 325 ms. We observed some sporadic transient responses to the onset of the holographic stimulation, although not systematically. Thanks to this temporal delay, these responses occurred in a temporal window that could be separated from the responses to the visual stimulus.

Since the AII amacrine cells targeted by holographic stimulation remained hyperpolarized during visual stimulation, we could assess the effect of hyperpolarization by comparing responses to visual stimulation alone with those elicited by combined visual and holographic stimulation (Fig. 4C). In many cases, we observed that the response to the visual disc was reduced when paired with holographic stimulation, compared to visual stimulation alone (Fig. 4C). Across the population of recorded OFF ganglion cells (*N* = 2 wild-type retinas, *n* = 50 cells), 23% of cells that exhibited a clear response to ON surround stimulation showed a significant decrease in this response during holographic stimulation (Fig. 4D, see Methods). We observed that the inhibition of AII amacrine cells located 5 RF radii away from a ganglion cell could produce a decrease of firing rate in surround responses between 20 Hz and 40 Hz. These results show that the RBC-AII amacrine cell pathway mediates the ON surround responses of OFF ganglion cells.

An alternative explanation for this decrease in response to ON surround stimulation is that the initial holographic stimulation desensitizes the photoreceptors, reducing their responsiveness to the visual stimulus presented 175 ms later. In that case, the decrease in response would be independent of AII amacrine cell activity. To test this hypothesis, we conducted control experiments using the same protocol but without gtACR1 expression in AII amacrine cells. In this condition, holographic stimulation impacts photoreceptors as before, but AII amacrine cells are not hyperpolarized. In these experiments, we observed no difference between the response to the visual stimulus alone and the combined holographic-visual stimulation (*N* = 2 wild-type retinas, *n* = 46 cells, Fig. 4C). Together, these results show that hyperpolarization of AII amacrine cells reduces the antagonistic surround modulation in OFF ganglion cells.

## Discussion

Using a combination of holographic stimulation and multi-electrode array recordings, we demonstrated that RBC activation can evoke responses in distant OFF ganglion cells, well beyond their classical receptive field center. We also found that hyperpolarizing AII amacrine cells decreases the response to ON surround stimulation in OFF ganglion cells. Our results thus show that the RBC-AII amacrine cell pathway participates in mediating the antagonistic surround modulation observed in OFF ganglion cells.

How can AII amacrine cell’s activation drive responses in distant OFF ganglion cells, given that AII amacrine cells are narrow-field and thus cannot form a direct synapse with these RGCs? A potential mechanism, also proposed by Lauritzen et al., (2019) based on anatomical data, suggests that AII amacrine cells could inhibit OFF cone bipolar cells, thereby reducing their excitatory input to OFF GABAergic wide-field amacrine cells. Alternatively, AII amacrine cells could inhibit OFF GABAergic wide-field amacrine cells directly (Fig. 5). In both cases, this could lead to a disinhibition of distant OFF cone bipolar cells, allowing them to excite OFF ganglion cells (figure 5). Wide-field GABAergic amacrine cells are involved in surround suppression for some types of ganglion cells (Baccus et al., 2008; Farrow et al., 2013; Nath et al., 2023; Park et al., 2020). Our results do not suggest a single circuit mediating antagonistic surround modulation in OFF ganglion cells. AII amacrine cells either directly or indirectly inhibit several types of wide-field GABAergic amacrine cells that are involved in surround suppression for different types of OFF ganglion cells.

**Fig. 5.**
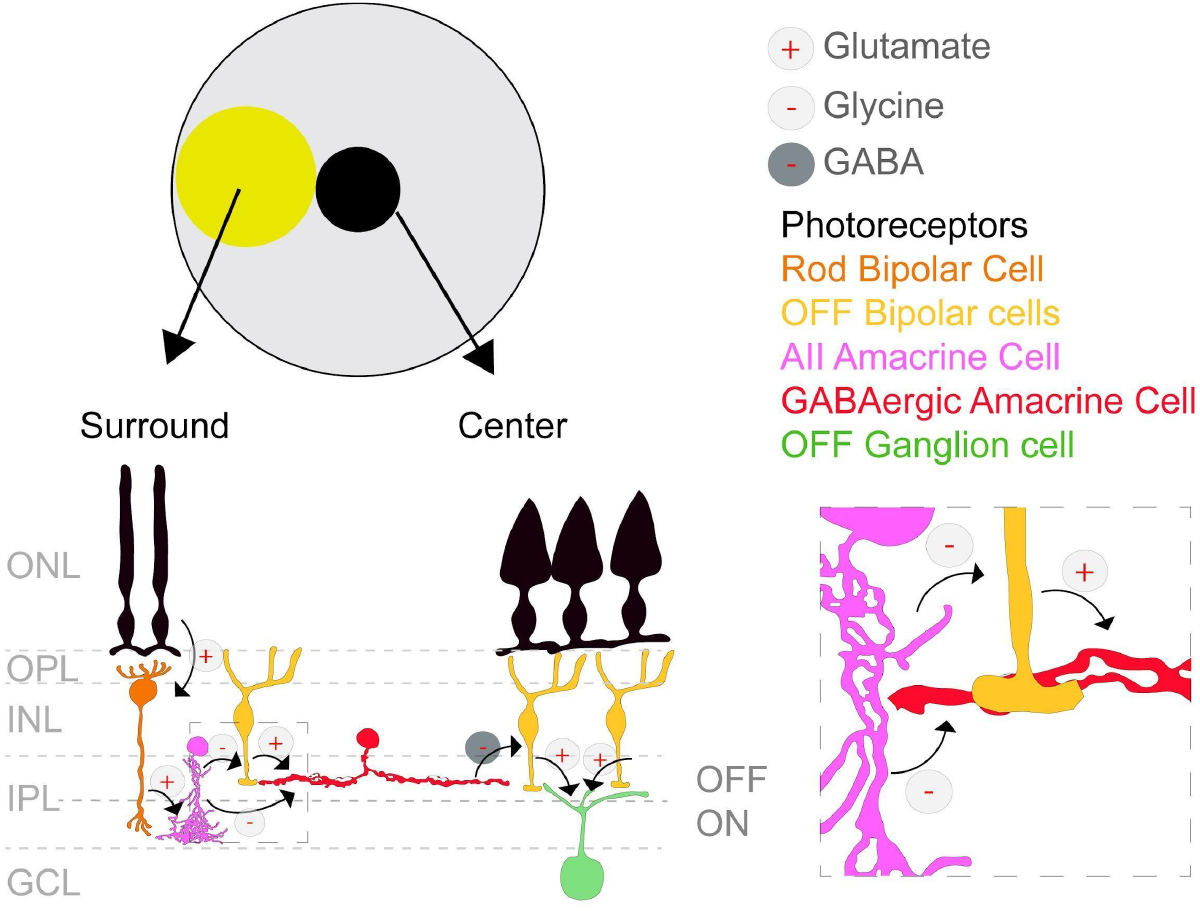
Scheme for the Role of the Rod Pathway in the Surround Response of OFF Ganglion Cells. Rod photoreceptors in the surround of OFF ganglion cells, hyperpolarized by a bright disc, activate rod bipolar cells. These rod bipolar cells, in turn, excite AII amacrine cells via glutamatergic transmission. The AII amacrine cells then modulate OFF ganglion cell activity via a disinhibitory circuit through two possible mechanisms: **1)** AII amacrine cells inhibit OFF bipolar cells, reducing their excitatory input to a GABAergic amacrine cell. This inhibition of the GABAergic amacrine cell ultimately increases the firing rate of the OFF ganglion cell. **2)** AII amacrine cells directly inhibit the GABAergic amacrine cell, leading to the same increase in OFF ganglion cell activity.

While our results demonstrate that AII amacrine cells participate in this modulation, it is likely that other pathways can also mediate a similar inhibition. Any glycinergic amacrine cell mediating crossover inhibition from the ON to the OFF pathway could also play the same role as the AII amacrine cell in the circuit described above. A more general mechanism to mediate antagonistic surround modulation in OFF ganglion cells could be that ON stimulation evokes a local crossover inhibition of the OFF pathway through ON bipolar cells and ON glycinergic amacrine cells. This inhibits the wide-field amacrine cells that mediate surround suppression for distant OFF ganglion cells. As a result, distant OFF ganglion cells can be disinhibited upon distant ON stimulation. In this more general circuit, the RBC-AII pathway is just one way to perform crossover inhibition, but not the only one. This suggests a mechanism where antagonistic surround modulation is implemented by having crossover inhibition piggyback the wide-field amacrine cells that mediate surround suppression.

Beyond these glycinergic mechanisms, additional pathways may also shape antagonistic surround modulation. In particular, lateral inhibition via horizontal cells and other amacrine cell networks can play an important role, and their relative contributions likely depend on factors such as background luminance and stimulus shape. Previous studies (Joesch and Meister, 2016; Szatko et al., 2020; Szikra et al., 2014) have shown that rods contribute to surround modulation, particularly through surround suppression, by laterally inhibiting cone output via horizontal cells. Our findings extend this understanding by showing that the pathway formed by RBCs and AII amacrine cells also plays a role in antagonistic surround responses in OFF ganglion cells. Previous studies (Münch et al., 2009; Demb and Singer, 2014) have also shown that, beyond relaying rod signals, AII amacrine cells can substantially reshape ganglion cell responses. Our study further highlights their role in generating antagonistic surround responses in OFF ganglion cells, leveraging their ability to respond to ON stimuli and inhibit OFF cells.

The RBC-AII pathway is critical for propagating rod signals in dim light. More recently, several studies (Ke et al., 2014; Pang et al., 2010; Szikra et al., 2014) have shown that this pathway remains active under mesopic light levels. This can be attributed to two factors. First, rods can transmit signals across a broad range of light levels before becoming saturated (Grimes et al 2018). Second, cones have been shown to connect to RBCs (Behrens et al., 2016; Pang et al., 2018). Our findings suggest that, under these conditions, the RBC-AII pathway contributes to distant surround modulation in OFF ganglion cells. It is also possible that AII amacrine cells have a more general role in shaping antagonistic surround responses, even beyond the light level where they receive inputs mostly from RBC. At higher light levels, AII amacrine cell activation through cone pathways (*e*.*g*., by receiving cone input (Pang et al., 2018) or ON bipolar cell input through gap junctions) could make them contribute to antagonistic surround modulation for a broader range of light levels, although the kinetics should be different if they are mostly driven by cones.

Finally, one of the most fascinating features of the brain is its ability to perform complex computations despite limited resources. The entire energy consumption of the brain is around 20 W, way below the energy required to run a supercomputer. Many studies have shown that this feat is achieved by saving resources. Efficient coding theories usually assume that the metabolic cost of firing is minimized (Balasubramanian and Berry, 2002; Niven, 2016). Some neural properties could be derived by minimizing the wiring length (Wang and Clandinin, 2016). More recently, using the connectome of the fly visual system as a constraint in a model allowed predicting many properties of the cell types of the fly visual system (Lappalainen et al., 2024). This suggests that wiring constraint is a key driver of the organization of the visual system.

If wiring constraints and the number of cells are precious resources that must be used efficiently, an effective strategy is to reuse existing elements whenever possible. In this context, individual cell types or circuits can be recruited to serve multiple computational roles across different situations. Our study illustrates this principle by showing how a single circuit, the RBC-AII pathway, is reused to perform distinct computations in different contexts. Similar forms of circuit reuse likely occur for other cell types, both within the retina and in other brain regions.

## Methods

### AAV Productions

Recombinant AAVs were produced by the triple transfection method (Grieger et al., 2016), and the resulting lysates were purified via iodixanol gradient ultracentrifugation as previously described (Dalkara et al., 2013). Briefly, a 40% iodixanol fraction was concentrated, and the buffer was exchanged using Amicon Ultra-15 Centrifugal Filter Units (Millipore, Molsheim, France). Vector stocks were then tittered for DNase-resistant vector genomes by real-time PCR relative to a standard (Aurnhammer et al., 2012). For experiments aimed at expressing CoChR (Klapoetke et al., 2014; Shemesh et al., 2018) in rod bipolar cells, we used the In4s-In3-200En-mGluR500P promoter (Lu et al., 2016), which has been previously demonstrated to enable specific expression of optogenetic proteins in these cells. To deliver the genetic material across the retinal layers, we used the AAV2-7m8 vector, a modified variant of AAV2 (Dalkara et al., 2013). For experiments targeting AII amacrine cells, we expressed the hyperpolarizing optogenetic protein gtACR1 (Govorunova et al., 2015) using the HKamac promoter, which has been shown to drive specific expression in AII amacrine cells (Khabou et al., 2023). The same AAV2-7m8 viral vector variant was also used.

### Animals And Intravitreal Injections

All experiments were done following Directive 2010/63/EU of the European Parliament. The protocol was approved by the Local Animal Ethics Committee of Paris 5 (CEEA 34). All mice used in this study were wild-type mice from Janvier Laboratories (Le Genest Saint Isle, France). For injections, mice were anesthetized with isoflurane (5% induction, 2% during the procedure). Pupils were dilated, and an ultrafine 30-gauge disposable needle was passed through the sclera, at the equator and next to the limbus, into the vitreous cavity. Injection of 1 µl stock containing 3.04 × 10^12^ vg/ml of AAV was made.

### Setup description

The optical setup was built around a commercial inverted microscope (Olympus, IX71). A 252-channel preamplifier (MultiChannel Systems) was placed on the stage of the microscope, and an MEA was aligned on top of the objective. All the optical paths passed through the inverted objective and the MEA. Three different optical paths were combined with different goals:

- 1p wide-field epifluorescence imaging. Imaging was obtained by collecting a 1P-induced fluorescence signal on a CCD camera (ORCA - 03G, Hamamatsu). Illumination was provided via a filtered light guide-coupled illumination system (UHGLGPS, Olympus).
- For visual stimulations, we focused the filtered (530/50 nm) light of a lamp (X-Cite Lumen Dynamics) on a digital micromirror device DMD (VIALUX LTD, 1024x768), which was generating the visual stimuli. Light from the DMD was projected through the objective (10x, NA 0.25, Olympus) and focused at the level of the photoreceptors.
- 2p optogenetic activation was achieved using computer-generated digital holography thoroughly described in (Ronzitti et al., 2017). Briefly, a femtosecond pulsed beam (InSight DeepSee, Spectra-Physics, fixed laser line 1040 nm) was expanded to illuminate the display of a spatial light modulator (SLM, LSH0700963, Hamamatsu). The SLM display was imaged on the back focal plane of the objective lens. The SLM was addressed with a phase mask calculated with custom-designed software (WavefrontDesigner) based on Gerchberg-Saxton iterative algorithms to produce arbitrarily defined intensity profiles at the sample plane and target opsin-expressing bipolar or amacrine cells.

### Multi-Electrode Array

All recordings were performed with a multi-electrode array of size 450 µm by 450 µm (16 by 16 electrodes with 30 µm spacing), at a sampling rate of 20 kHz. The data was sorted to reconstruct the spiking activity of each ganglion cell using the software Spyking Circus (Yger et al, 2018). Artifacts due to the holographic stimulation were removed by high-pass filtering the extracellular signal (cut-off frequency: 200 Hz), and then removing the signal in 10 ms windows centered on the onsets and offsets of the stimulation.

### Light Stimulation

Visual stimulation was delivered through a Digital Micromirror Device (DMD). To estimate ganglion cell receptive fields we used a white noise stimulus (checkerboard). Each frame consisted of a grid of gray checks of size 50µm each, arranged in an array of size 38 by 51. The light intensity of each check changed randomly at each frame according to a normal distribution. This stimulus was played at a 30 hz rate for a total time of 35 to 45 minutes, depending on the experiment.

### Holographic Stimulation

One-photon widefield fluorescence imaging was used to image the bipolar cell layer and identify the RBCs expressing the opsin. We chose a subset of RBCs and determined their position to deliver the optogenetic stimulus. For optogenetic stimulation, we relied on cell-targeted two-photon computer generated holography.

For individual activation (Fig. 2), we identified several expressing RBCs. We targeted these cells one by one at maximum intensity for time intervals of 500 ms, interleaved with 1 s of pause. We collected 20 trials for each RBC activation, for a total protocol time of around 20 minutes, using a stimulation power of 0.09 mW/μm^2^.

For the control protocol (Fig. S1), the locations of the holographic patterns were chosen arbitrarily. We generated a grid of 49 holographic locations (7 by 7) equally spaced across the whole multi-electrode array. These locations were targeted one by one for a time interval of 500 ms, interleaved with 1 s pauses. For each pattern, we collected 20 trials, for a total time of around 25 minutes.

### Joint Light And Holographic Stimulation

For the AII amacrine cells inhibition protocol, we combined a visual stimulus with the holographic activation described above. The visual stimulus consisted of a white light disc of diameter 250 µm, displayed on one side of the multi-electrode array. Inside the perimeter of the disc, we selected 15 holographic locations, equally distanced with a spacing of around 45 µm (Fig. 4).

The holographic stimuli were delivered for a time interval of 500 ms each, interleaved with pauses of 3.5 s. For each pattern, we collected 20 or more trials. During the holographic stimulation, we displayed the white disc, after 175 ms from the onset of the holography, for a time interval of 325 ms. This delay between the start of visual and holographic stimuli was needed to avoid confounding between the responses of ganglion cells due to optogenetic activation by holography and the ones due to visual activation by the disc. We interleaved the composite stimulation with presentations of the visual stimulus alone, for a total time of 45 minutes.

### Receptive Fields

To estimate the ganglion cell receptive fields, we first calculated the spike-triggered averages (STAs) of the responses to the checkerboard stimulus. The averages were computed on the 21 checkerboard frames preceding a stimulus, for a total period of 700 ms. These averages can then be described with a three-dimensional matrix, where each value represents the average light intensity of a given check for a given frame preceding the stimulus.

The STAs were then factorized into spatial and temporal components. The spatial component is obtained by computing the standard deviation of the mean of the STA across frames. This produces a two-dimensional heat map matrix, where each value is an index representing how much the corresponding check produces a response in the ganglion cell. The corresponding receptive fields were obtained by fitting a two-dimensional Gaussian distribution to the spatial STAs. We generated the ellipse corresponding to the region for which the Gaussian distribution had a standard deviation equal to one and estimated the receptive field center of the ganglion cell as the surface delimited by this ellipse. The temporal component was then calculated by considering only the checks lying inside the receptive field centers and averaging their STA values across the two spatial dimensions. All the cells for which it was not possible to estimate a receptive field center were excluded from the study.

### Activation Score

To detect and quantify the ganglion cell responses produced by RBC and AII amacrine cell activation we defined an activation index *A* and an activation score *S*. We first computed an activation threshold *tresh*_*up*_ as follows: for a given stimulus *s* and ganglion cell *c*, we considered a control window of 300ms antecedent to stimulus onset and computed the average mean spontaneous firing rate 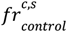 and its standard deviation 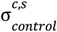 across all trials. We defined the activation threshold as:

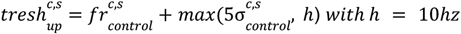

Then, we looked at a response window of 400ms after the presentation of the optogenetic stimulus (excluding respectively the first and last 50ms of stimulation) and computed the peri-stimulus time histogram *psth*^*c,s*^ (time bin equal to 50ms). We assigned an activation index *A*^*c,s*^ equal to 1 if the ganglion cell mean response *psth*^*c,s*^ exceeded its activation threshold 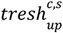 for at least one time bin, or a score equal to 0 otherwise. We then defined the activation score S as:

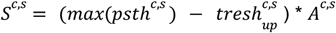

We then extended our previous definitions of activation index and activation score as follows:

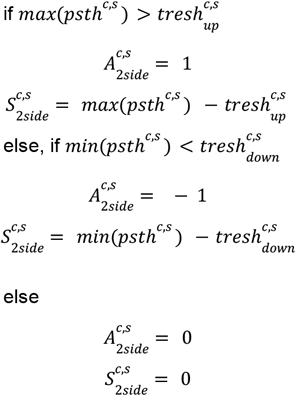

Positive activation index and scores correspond to a detected ganglion cell activation. Negative index and scores correspond to ganglion cell inhibition. Null index and scores entail no response was detected.

### Comparative Activation Score

To quantify the AII amacrine cells’ modulation of ganglion cell visual responses, we defined a comparative activation score. This score measures how much, in terms of firing rate, the AIIs inhibited by holographic stimulation contribute to the ganglion cell response. First, we looked at the ganglion cell responses to the pure visual stimulus. We considered a response window of 350 ms, starting 75 ms after stimulus onset, and computed the mean across trials. We followed the same procedure for the response to the composite visual and holographic stimulation. The comparative activation score is then defined as the maximum of the visual response minus the maximum of the visual and holographic response (Fig. 4D)

### Normalized Distances

Normalized distances between pairs of RBC and ganglion cells (RGC) were computed as the Euclidean distance on the imaging frame divided by the ganglion cells’ receptive field radius:

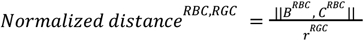

Where *B*^*RBC*^ is the position of the RBC on the imaging frame, *C*^*RGC*^ is the center of the ganglion cells receptive field center projected on the imaging frame, and *r*^*RGC*^ is the longest radius of the ganglion cells’ receptive field center. The same formula was used for normalized distances between ganglion cells and holographic patterns.

### Effects Of RBC Stimulation: Population Analysis

To quantify the effect of RBC stimulation on ganglion cells responses at different distances, we first computed the activation score of each ganglion cell for every RBC stimulation. For each RBC-ganglion cell pair, we calculated their normalized distance. Then we binned all the pairs according to their distances, considering a range of 0 to 5 normalized distances (bin size 0.33). For each distance bin, we computed the probability of observing a ganglion cell’s response to RBC stimulation as the number of pairs with activation scores different from zero, divided by the total number of pairs in the bin (Fig. 2. E, F).

Finally, we used the same analysis to assess the responses to holographic stimulation mediated by photoreceptors in the absence of blockers and opsin expression. In this case, we used the extended version of our activity index to compute our probability distributions (see section above), as we wanted to assess both the inhibitory and excitatory effects of the stimulation. As expected, the holographic stimulation had an excitatory effect on ON ganglion cells and an inhibitory effect on OFF ganglion cells. These effects were stronger and more frequently observed for holographic spots located inside the ganglion cell receptive field centers, and got progressively weaker for spots located further away (Fig. S1).

### Responses To AII Amacrine Cells Inhibition: Population Analysis

To estimate the contribution of AIIs to the responses of OFF ganglion cells, we computed the activation score of each ganglion cell to the pure white disc stimulus as described above and selected all the ganglion cells with a non-zero activation score. For each of these ganglion cells, we computed the relative distances to all the holographic locations. We binned all the ganglion cell-holographic pattern pairs according to their relative distance, considering a range of 0 to 10 normalized distances (bin size 0.66). For each distance bin, we calculated the average comparative activation scores (Fig. 4).

To quantify the extent of AII modulation of OFF ganglion cell’s surround responses, we selected only the ganglion cells for which the disc lies outside of their receptive field center (50 ganglion cells). We considered all pairs of these ganglion cells and holographic patterns and performed the same distance binning explained above. Then, we computed the probability of observing a surround modulation as follows: for each distance bin, we counted the number of ganglion cell-holographic spot pairs with a comparative activation index above 10 hz and divided by the total number of pairs (Fig. 3.D).

## Data analysis

For Figures 2 and 4, statistical differences were assessed using a one-sample t-test to determine whether the mean activation in each distance bin was significantly different from zero. Additionally, for Figure 2, a paired t-test was performed to compare activation z-scores between control conditions and after the application of strychnine. All statistical analyses were conducted using a custom MATLAB script. Data values are presented as mean ± standard error (Fig. 2 and 4) and as mean ± standard deviation (Fig. 3). Significance levels are indicated as follows: * p < 0.05; ** p < 0.01; *** p < 0.001.

## Author contributions

**G.S**. and **F.T**. conceived, designed, performed experiments, analyzed, interpreted the data, and reviewed the submitted version. **V.C.G**. data analysis, interpreted the data, figures preparation, wrote the original draft, reviewed and edited the submitted version. **T.B**. designed, interpreted the data, performed experiments, and reviewed the submitted version. **E.O**. and **B.S**. performed experiments and reviewed the submitted version. **D.D**. performed the AAV production and reviewed the submitted version.. **E.R**., **E.P**., and **V.E**. designed and built the holography path and reviewed the submitted version. **O.M**. Conceptualization, funding acquisition, project administration, supervision, visualization, wrote the original draft, and reviewed the submitted version.

## Supplementary information

**Fig S1.**
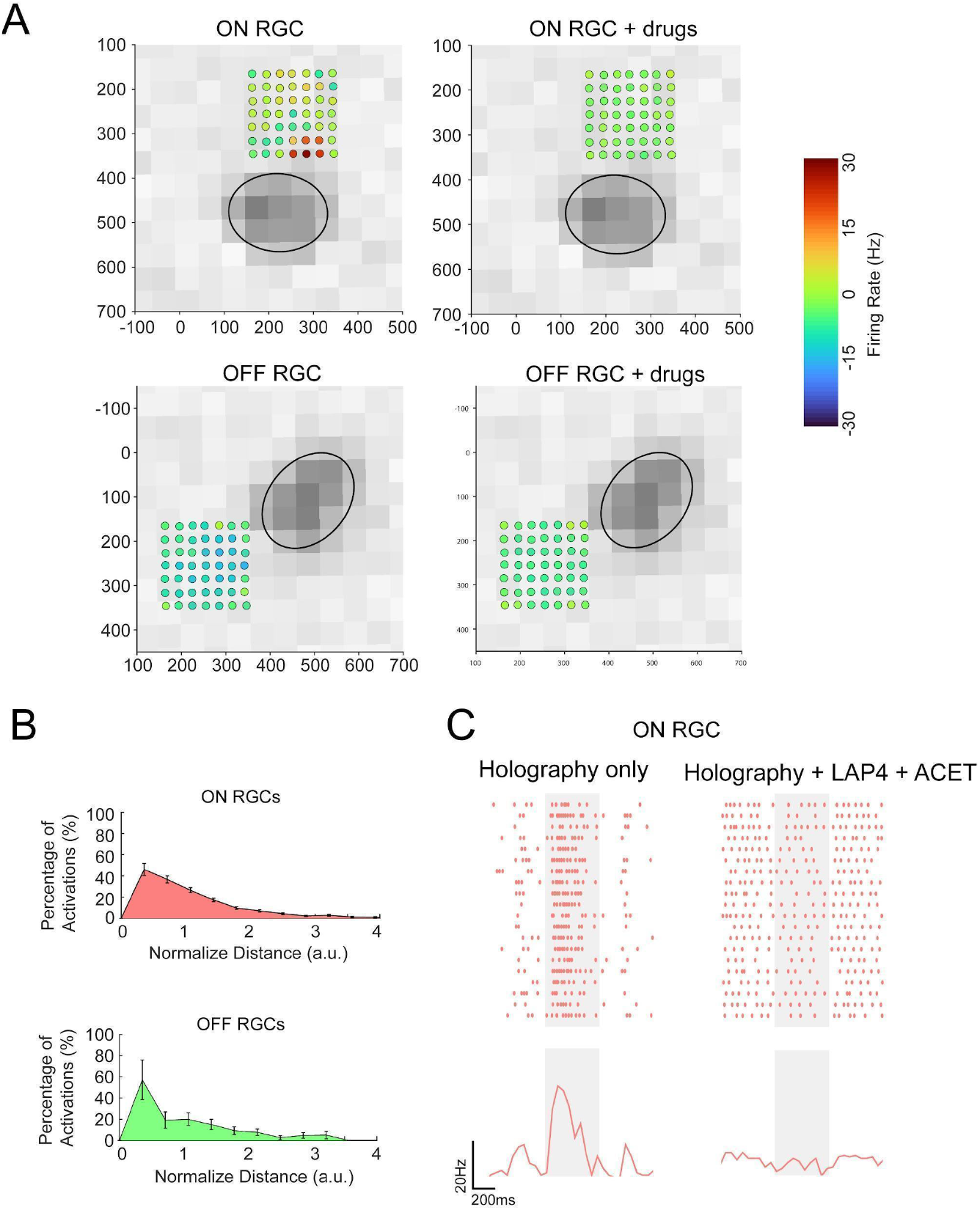
Pharmacological blockers prevent all ganglion cells’ visual responses to the holographic stimulation. **A**. responses of a representative ON ganglion cell to the holographic stimulation in the absence of blockers and with no opsin expressed. On the left: the spiking activity of the cell across different trials. On the right: the mean response. The time interval of the stimulation is depicted by the grey regions. **B**. responses of the same ON ganglion cell shown in A after the application of blockers **C**. left: Responses of the representative ON ganglion cell shown in A and B to all the holographic patterns in the absence of blockers and with no opsin expressed. The background images represent the spatial STAs. The black ellipse indicates the spatial extent of the receptive field center. The colored dots show all the locations targeted with the holographic stimulation. The colors of the dots represent the peak firing rates of the corresponding induced responses (legend on bottom). The holographic spot shown in A and B is marked with a yellow circle. Right: same as left after the application of blockers. **D**. same as C for a representative OFF ganglion cell.

